# Targeting RyR2 with a phosphorylation site-specific nanobody Reverses Dysfunction of Failing Cardiomyocytes in Rat

**DOI:** 10.1101/435875

**Authors:** Tian Li, Yafeng Shen, Fangxing Lin, Wenyan Fu, Shuowu Liu, Xiaoyan Fan, Xuting Ye, Ying Tang, Min Ding, Yongji Yang, Changhai Lei, Shi Hu

## Abstract

Chronic PKA phosphorylation of RyR2 has been shown to increased diastolic SR Ca2+ leak and lead to cardiac dysfunction. Since the change of phosphorylation level of RyR2 is a biomarker of failing heart, we attempted to verify the hypothesis that intracellular gene delivery of a RyR2 targeting phosphorylation site-specific nanobody could preserve contractility of failing myocardium. In present study, we acquired the RyR2-specific nanobodies from a phage display library which are variable domains of camellidae heavy chain-only antibodies (VHH). One of the monoclonal nanobodies, AR185, inhibiting RyR2 phosphorylation in an in vitro assay was then chosen for further investigation. We investigated the potential of adeno-associated virus (AAV)-9-mediated cardiac expression of AR185 against post-ischemic heart failure. Adeno-associated virus gene delivery elevated the intracellular expression AR185 protein in the ischemic heart failure model of rats, and this treatment normalized the systolic and diastolic dysfunction of the failing myocardium in vivo and in vitro by reversing myocardial Ca^2+^ handling. Furthermore, AR185 gene transfer to failing cardiomyocytes reduced the frequency of sarcoplasmic reticulum (SR) calcium leak, thereby restoring the attenuated intracellular calcium transients and SR calcium load. Moreover, AR185 gene transfer inhibited PKA phosphorylation of RyR2 in failing cardiomyocytes. Our results provided strong pre-clinical experimental evidence of the cardiac expression of RyR2 nanobody with AAV9 vectors as a promising therapeutic strategy for ischemic heart failure.

## Introduction

In most countries, heart problems are the leading cause of morbidity and mortality(*1*). Heart failure is the end stage of most heart diseases, with approximately 38 million heart failure patients worldwide(*2*). Despite the use of drugs, implanted cardiac assist devices and surgical treatments, many patients’ conditions will still irreversibly deteriorate, eventually being difficult to control and rescue.(*3*) Therefore, studying the molecular mechanism of cardiovascular pathology, developing new therapeutic strategies and preparations have the great significance for the prevention and treatment of heart failure. No matter what the cause, heart failure patients in the late stage of the disease have a common feature that calcium circulation is abnormal in the cardiomyocytes.! A hallmark of failing cardiomyocytes is the change of excitatory contraction coupling, including the reduced amplitude of Ca2+ transients, delayed onset and decay kinetics of Ca2+ transients. These changes eventually lead to decreased contractility, delayed contraction, and reduced diastole. In addition, a rare spontaneous Ca2+ release events (calcium leaks) in the resting period of healthy cardiomyocytes occurs more frequently. All the change of Ca2+ handling is attributed to impaired function of RyR2, SERCA2a, Na+-Ca2+ exchanger (NCX).! Changes in expression and function of these ion channel proteins and their negative effect after MI have been reported in many studies (*4, 5*), which directly attenuate the cardiac contractility. Therefore, preventing these detrimental effects of myocardial infarction, as well as the effects of ischemic injury due to the changes in expression or function of calcium-channel proteins, is an optimal treatment strategy for cardio-protection.

The activity of RyR2 function is regulated by various mechanisms, and many factors will affect it. The post-translational regulation abnormality of RyR2 is the main cause of the dysfunction of RyR2 protein. Phosphorylation is a crucial post-translational modification of RyR2 protein.(*6, 7*) The "calcium leakage" caused by RyR2 hyperphosphorylation is considered to be an important pathological mechanism for myocardial injury and heart failure development. (*8, 9*)The study found that the expression of catecholamines in the blood of chronic heart failure patients was up-regulated, and intracellular protein kinase A (PKA) levels and activities continued to increase. After PKA is continuously activated, hyperphosphorylation of RyR2 protein causes an increase of the dissociation rate of FKBP12.6 and RyR2, resulting in a change of RyR2 structure, which increases the sensitivity of RyR2 to calcium ions, resulting in a small amount of calcium ions can stimulate the release of RyR2 channels, causing the calcium leakage in the resting stage of cardiomyocytes. The earliest article demonstrating changes in ryr2 phosphorylation in patients with heart failure was published by Marx et al. in 2000(*10*). Marx et al.(*10*), Wehrens et al(*11*). and Shan et al.(*12*) proposed a hypothesis: sympathetic excitation in heart failure, activation of PKA-mediated RyR2 S2808 Hyperphosphorylation, while inhibiting the binding of FKBP12.6 to RyR2, increases the probability of RyR2 opening.

But this assumption is currently controversial. The biggest controversy is whether RyR2 S2808 hyperphosphorylation plays a decisive role in heart failure. Wehrens et al. (*11*)believe that RyR2 S2808 phosphorylation plays a critical role in the experimental myocardial ischemia-induced heart impaired process. They established a heart failure model which induced by myocardial infarction in RyR2 S2808 knockout mice and wild-type mice. Heart function tests were performed 4 weeks after ischemia. The results showed that RyR2 S2808 knockout mice showed significant improvement in cardiac function such as ejection fraction, shortening fraction, maximal rate of left ventricular pressure. At the same time, they also found that RyR2 of RyR2 S2808 knockout mice could not be phosphorylated by PKA, and PKA lost regulation of FKBP12.6, resulting in FKBP12.6 not being able to dissociate from RyR2. However, Zhang et al.(*13*) and Houser et al.(*14*) believe that PKA-mediated hyperphosphorylation of RyR2 S2808 does not alter myocardial contractility and does not improve symptoms of heart failure and arrhythmia. They used a similar experimental protocol and found no improvement in the heart function of the mouse knocked out by RyR2 S2808. However the biggest problem with Hauser et al. is that the number of samples is too small. Inhibition of RyR2 phosphorylation may be one of the effective treatment of ischemic heart failure.

Here, we hypothesis that targeting RyR2 using anti-phosphorylation agents may improve treatment efficacy. We identified a camel single-domain antibody to RyR2 that have the ability to inhibit PKA dependent S2808 phosphorylation in vitro. To evaluate its potential effect in the treatment of heart failure, an adeno-associated virus (AAV) based intracellular antibody delivery strategy were adopt to achieve cardiac-specific gene-therapy and demonstrated therapeutic effect both in cell based assays and in vivo models.

## Materials and Methods

### Cell Culture

1-2-day old Sprague-Dawley neonatal rats were used to isolate the neonatal rat ventricular myocytes. Neonatal rats were sterilized with alcohol and sacrificed by decapitation. Then opening chest in asepsis condition, whole hearts were removed. Ventricles were cut from whole hearts and incubated overnight in 1x Hank’s Buffered Saline Solution (HBSS) with 0.06% trypsin at 4°C. The next day, the reagent that processed cardiomyocytes was converted into 1x HBSS with 10mg/mL Collagenase II (Worthington Biochemical), cardiomyocytes were isolated into individuals. To remove contaminating fibroblasts, we pre-plated the resulting suspensions of cardiomyocytes, and a density of 1 × 10^6^ cells/well was plated in 6-well dishes which was cultured with Dulbecco’s Modified Eagle Medium (DMEM, Invitrogen) containing 10% fetal bovine serum. Before seeding, 0.1% gelatin was used to pre-treat all tissue culture plates that were inoculated with cardiomyocytes for at least 1 h. After 24 h incubation, 1x PBS was used as rinse to wash the cardiomyocytes and myocytes were cultured in DMEM with 0.5x Nutridoma-SP (Roche Applied Sciences).

### Purification of RyR2 by GST-FKBP12

The expression and purification of GST-FKBP12 were reported previously(*15, 16*). Briefly, pGEX-4T-2 vector carried the cDNAs encoding rat FKBP12 with a C-terminal 6×His tag. To overexpress the protein, plasmids were transformed into BL21 (DE3) strain. When the OD600 of BL21 reached 1.0, adding 0.4 mM IPTG for induction overnight and the temperature was adjusted to 16°C. Cells were collected, lysed and centrifuged, Ni2+-NTA resin (Qiagen) was used to purify the recombinant protein and eluted by imidazole containing buffer. The protein was subjected to Anion exchange chromatography (SOURCE 15Q, GE Healthcare) for fine fractionation.

To prepare the sarcoplasmic reticulum membrane of cardiomyocytes, a single rat heart was cutted into small pieces, and resuspended in 5 volumes of homogenization buffer that formulated with 25 mM Tris, pH 7.5, 150 mM NaCl, 5mM EDTA, and the protease inhibitor cocktail (0.2 mM PMSF, 1.3 μg/ml aprotinin, 0.7 μg/ml pepstatin, and 5 μg/ml leupeptin). The blender was used to homogenize samples fifteen cycles. Low speed centrifugation (6,000 g) was performed to remove the debris for 6 min. After 1 hour high speed centrifugation (20,000g), the precipitation in the supernatant obtained by the last step was preserved and resuspended in 2 volumes of homogenization solution. Liquid nitrogen was used to quickly freeze the suspension

In this study the same strategy for the purification of RyR1(*15*)and the RyR2 (*16*) were adopted to purify RyR2 protein. The homogenization buffer plus 2% CHAPS, 1% soybean lecithin, and 2 mM DTT dissolved the sarcoplasmic reticulum membrane of cardiomyocytes (a quarter of total membrane from a single heart) at 4°C for 2 hours in. The whole system extracted approximately 10 mg of GST-FKBP12. Precipitate was discarded by ultra-high-speed centrifugation (200,000 g) and the supernatant was applied to GS4B column (GE Healthcare) for affinity purification. The washing buffer of resin was similar to the homogenization buffer, except that NaCl concentration was adjusted to 1M, 2mM DTT and 0.1% Digitonin were added in the buffer. The complex was eluted by elution containing 75 mM Tris-HCl, pH 8.0, 150 mM NaCl, 10 mM GSH, 0.1% Digitonin, 2 mM DTT, 5 mM EDTA, and protease inhibitors. The eluted protein was finally purified by size exclusion chromatography (SEC, Superose 6 10/300 GL, GE Healthcare) in the buffer containing 25 mM Tris, pH 7.5, 300 mM NaCl, 0.1% Digitonin, 2 mM DTT, 5 mM EDTA, and protease inhibitors.

### Phage Display and Biopanning

Camel library construction were reported previously (*17, 18*). Peripheral blood mononuclear cells (PBMCs) were isolated from a total of 300ml blood sample, taken via Leucosep^®^ tubes (Greiner Bio-One, Frickenhausen, Germany). According to previous report, total RNA was extracted and nested PCR cloned VHH genes (*19*). Phagemid vector pCANTAB5E (GE Healthcare Life Science, Pittsburgh, USA) carried the final PCR products (~ 400bp) and introduced into electro-competent E. coli TG1 cells that was freshly prepared. Cells were selected on LB agar plates supplemented with ampicillin and glucose cultured overnight at 37°C. After scraped from the plates, the colonies were stored at −80°C in LB supplemented with 20% glycerol.

Repetitious biopanning was performed to enrich the phage clones as described previously (*20*). 100 μg/ml RyR2 was covered on the 96-well Maxisorp plate overnight at 4°C. Both the plate and 10^11^−10^12^ pfu of phage were blocked with 1% skimmed milk for 1 hour at room temperature. Then pre-blocked phage supernatant was added to each well to allow binding. After 1 hour of incubation at room temperature, the unbound and nonspecifically bound phages were removed using 5 washes. 100 μl pH 2.0 elution buffer eluted the specifically bound phage with incubating 10 minutes at room temperature. 30 μl of 1 M Tris-HCl buffer (pH 8.5) neutralized the eluate and the eluate was used to infect freshly prepared E. coli TG1 cells. After four rounds of panning, 300 randomly picked clones were analyzed for RyR2 binding by phage ELISA.

### ELISA

The phage ELISA was performed as previously described(*21*). Briefly, 50 μl of 5 μg/ml RyR2 was covered on Nunc MaxiSorp 96-well flat-bottomed plates overnight at 4°C. Both the plate and phage were blocked with 1% skimmed milk for 1 hour at room temperature. Pre-blocked phage supernatant was then added to the plate. The antibody that was a horseradish peroxidase (HRP)-conjugated mouse anti-M13 antibody (GE Healthcare) was used to detected the binding property.

### Expression and purification of anti-RyR2 nanobody fragments

Nanobody-encoding gene segments were re-cloned in vector pUR5850. Recombinant nanobody fragments were expressed in the periplasm of E. coli and purified by means of immobilized metal ion affinity chromatography (IMAC). Both experiment were performed as previously described (*17*).

### RyR2 Phosphorylation

RyR2 was incubated with 500 μg rat ventricle homogenate and anti-RyR antibody in 0.5 ml modified radioimmunoprecipitation assay buffer(Ripa; 50 mM Tris HCl buffer (pH 7.4) / 0.9% NaCl / 5.0mM NaF / 1.0mM Na_3_VO_4_ / 0.5% Triton X-100/protease inhibitor) to be immunoprecipitated as described (*11, 22*). Samples were incubation with protein A-Sepharose beads for 1h and then transferred into 10 μl of 1.5× phosphorylation buffer (8 mM MgCl_2_/10 mM EGTA/50 mM Tris/Pipes, pH 6.8) added either PKA catalytic subunit (Sigma, St.Louis, MO) or inhibitor PKI5–24 (500 nM, Calbiochem, San Diego) with or without different nanobody fragments (10 ug/ml). For maximally dephosphorylating PKA sites, alkaline phosphatase (AP) was used to pretreated samples before PKA phosphorylation. [^32^P]-ATP standards were used to determine the stoichiometry of phosphorylation of RyR2 by PKA. To calculate he [^32^P]/RyR2 ratio, the amount of high-affinity [^3^H]-ryanodine binding dividded [^32^P]-phosphorylation of RyR2 which was described previously(*11, 22*).

### AAV9 vector production

To express an upstream eGFP reporter, the construct of the intracellular antibody fragment, AR185 or AR117, was engineered and a Self-cleaving 2A peptide (T2A) separated the reporter from the nanobody sequence. The sequence that encoded the construct of whole fragments was subcloned into the pAAV vector(*23*) resulting in AAV.AR185 and AAV.AR177. To produce AAV9 pseudotyped vectors, there were three plasmids cotransfected in 293T cells, shuttle plasmid carrying the transgene flanked by inverted terminal repeats (ITRs), pAAV-RC9 plasmid encoding the AAV-9 rep and cap sequence, and pHelper plasmid containing the adenovirus genes that were necessary for AAV reproduction. There was a set of acknowledge procedures for production, purification, and titration of AAV vectors that was described (*24–27*).

### Experimental rat HF model

All animal procedures and experiments were performed with the approval of the animal ethics committee of the Second Military Medical University. The animal experiments were performed according to the NIH guidelines (Guide for the care and use of laboratory animals). Either myocardial infarction (MI) (*28*) or a sham procedure were randomized to performed on ten to twelve-week-old Sprague-Dawley male rats. Briefly, after deep anesthesia, rats were fixed on the operating table and faded. After intubation and placement on a respirator, a left lateral thoracotomy was performed between the 3rd and 4th ribs where closed the left edge of the sternum, and a ligature was placed 2 mm below the origin of left anterior descending coronary artery. Ischemia was verified by electrocardiogram, and then the chest was closed. After resuming spontaneous breathing, the rat was extubated and returned to its cage. Sham-operated animals were subjected the same procedure except for the absence of coronary artery ligation(*28*). The ligated rats were further randomly divided into three experimental groups as follows: heart failure group, AAV AR185 treated (AAV.AR185) group and AAV AR117 treated (AAV.AR117). A sterile syringe and 29-gauge needle were used to inject 300 μL bolus of AAV vector (treated groups) or saline (heart failure or sham group) into the tail vein of the rat.

### Ultrasound cardiographs

Eight weeks after tail vein injection, the cardiac function of the rat was examined by a Visual Sonics Ultrasound system. Rats were anesthetized with isoflurane. Then Direct datum of left ventricular contraction such as left ventricular end-systolic diameter (LVEDs) and left ventricular end-diastolic diameter (LVEDd) was recorded and calculated as specific evaluation indicators and analyzed such as fractional shortening (FS) and ejection fraction(EF).

### Transmission electron microscopy

Eight weeks after tail vein injection, paraformaldehyde and osmium tetroxide were respectively used to fix the fragments of left ventricular tissue in two steps. The fragments were dehydrated through a graded ethanol and acetone series and then embedded in Epon. LEICA EM UC6 (Leica, Austria) was used to cut ultrathin sections of fragments in Epon that were subsequently stained with uranyl acetate and lead citrate. The electron microscopy samples of myocardial tissues were observed by H-7650 transmission electronic microscope (TEM, Hitachi, Japan) and ultrastructural images of tissues were analyzed.

### Single ventricular myocytes isolation, laser scanning confocal microscope analysis and cardiac myocyte contractility

The cardiomyocytes were isolated from left ventricles by enzymatic dissociation method. In brief, rats were anesthetized with pentobarbital sodium (40 mg/kg, intraperitoneal), the hearts were excised and perfused by Tyrode’s solution(in mM: 135 NaCl, 5.4 KCl, 0.33 NaH_2_PO_4_, 10 glucose and 5 HEPES, pH was adjusted to 7.30 with NaOH) with or without collagenase type ‖ at a rate of 8 ml/min in a Langendorff system after immediate aortic cannulation . Hearts were firstly perfused ©, eight minutes later, the Tyrode’s solution was replaced by Tyrode’s solution containing 0.64 mg/ml collagenase type ‖‖ (335 U/mg), 0.2% BSA and 57.6 μM CaCl. Collagenase digested the heart continuously until the cardiac apex became soft. Left ventricular tissue was then cut into pieces in Kraft-Bruhe solution (in mM: 50 L-glutamic acid, 30 KCl, 80 KOH, 30 NaH_2_PO_4_, 20 taurine, 10 HEPES, 10 glucose, 3 MgSO_4_ and 0.47 EGTA, pH was adjusted to 7.35 with KOH) and mechanical method was further performed to obtain single ventricular cardiomyocytes. In the experiment, both of the Tyrode’s and Kraft-Bruhe solution were pre-warmed to 37°C and bubbled with 100% O_2_, and all these experiments were conducted at room temperature (22-24°C).

Leica TCS SP2 confocal microscope (Leica, Germany) was used to detected the fluorescence that represented intra-cellular and intra-SR Ca^2+^ concentration in xyt and xt modes and exported the images of intra-SR Ca^2+^ content, Ca^2+^ transient and Ca^2+^ sparks. After 30 min incubation with 10 μM Fluo-4AM (purchased from East-Chemical Technology Co., Ltd.) at 37°C, the intra-cellular calcium transients induced by caffeine and Ca^2+^ sparks in isolated cardiomyocytes were measured. Other isolated cardiomyocytes were loaded with 10 μM Fluo-5N (purchased from East-Chemical Technology Co., Ltd.) for 3 h at 37°C and then the intra-SR Ca^2+^ content was measured. The 488-nm line of an argon-ion laser was the exciting light of Fluo-4AM and Fluo-5N, and the fluorescence was detected at wavelengths of 500–560 nm.

For quantitative analysis, the temporal dynamics in fluorescence were expressed as Δ*F*/*F*_0_ = (*F*–*F*_0_)/*F_0_*, where *F* represents fluorescence intensity at time t and *F*_0_ stands for baseline intensity. Cardiomyocytes were placed in a chamber that was mounted on the stage of an inverted microscope (Zeiss X-40; Go’ttingen, Germany) with Tyrode’s solution containing 1.2mM CaCl at room temperature (22–24°C). Field stimulated (10 V) at a frequency of 1 Hz and pulse width of 10 ms was applied to cardiomyocytes by a pair of platinum electrode. Relevant data of contractile kinetics of the myocytes were detected by high-speed video edge-detection system (IonOptix, Milton, MA)

### Immunoblotting

Dead cells were eliminate by rinsing with PBS and then the cells were homogenized in cold lysis buffer containing (in mM: 75 Tris–HCl, 225 NaCl, 1.5 EDTA, Nonidet P-40 (4.5%, w/v), 5 sodium vanadate, 40 sodium fluoride, 10 sodium pyrophosphate, 10 N-ethylmaleimide and a protease inhibitor cocktail (Complete, Roche Diagnostics, Mannheim, Germany), pH adjusted to 7.4). The supernatant was collected after 10 min centrifugation (10,000 g) of lysate at 4°C. Proteins were separated and analyzed by SDS-PAGE (3.5–8% gradient gels), transferred to PVDF membrane, and immunoblotted with indicated antibodies. All the assays were repeated three times and the results shown are from one representative assay.

### Tissue analysis

Tissue sections were processed into slides either after being formalin-fixed and paraffin-embedded (FFPE) or embedded in non-fixed optimal cutting temperature compound (OCT, Tissue-Tek); tissue samples were also lysed in the CelLytic MT Lysis reagent (Sigma). Immunohistochemistry slides were scanned using an Imagescope (Scan-Scope AT, Aperio). The We defined the sum of the values obtained by multiplying the staining intensity and proportion as H-score that ranged from 0 to 300. Ratios of pRyR2 to total RyR2 were determined by western blot and ELISA.

### Statistical analysis

Statistical analysis was performed using Student’s unpaired t-test to identify significant differences unless otherwise indicated. Multiple comparisons were conducted using a two-way ANOVA followed by the Bonferroni post hoc test; a P-value less than 0.05 was considered a significant difference.

## 3 Results

### Generation of Anti-RyR2 nanobodies that specifically inhibits the Phosphorylation of RyR2 S2808

We first obtained and purified RyR2 from rat heart by using GST-fused FKBP12 as the published strategies described. To construct the camel VHH library, blood samples of 30 non-immunized, four year-old male Bactrian camel were collected. B lymphocyte cDNA encoding VHHs was used to construct a phage display VHH library that consisted of approximately 3 × 10^8^ individual colonies. VHH gene corresponded to the size of insert of over 98% colonies. For confirming the heterogeneity of the individual clones from the library, we sequenced fifty randomly selected clones, and each clone showed a distinct VHH sequence.

In order to select nanobodies with specific ability to bind RyR2, bio-panning was performed with immobilized RyR2 protein. After the third round of panning, the result showed an obvious enrichment of phage particles that carried RyR2-specific VHH (Fig. 1A). Phage clones exhibited increased binding to RyR2 after the second round of panning. During four rounds of panning there was no phage clone that was found binding to BSA (Fig. 1B). VHH fragments of 300 individual colonies that were randomly chosen were expressed in an ELISA for screening colonies which bound to RyR2. Among these clones, 276 antibody fragments specifically bound to RyR2. One antibody fragment which did not bind to RyR2 was choose as a negative control, termed as VHH-AR117. To obtain antibodies that functionally inhibit of RyR2 phosphorylation, each of the antibody fragments was tested for its effect in an ELSA based RyR2 phosphorylation assay. 4 antibody fragments were potent inhibitors of RyR2 phosphorylation. The complementary determining regions (CDRs) were confirmed by sequence analysis and the result revealed that there was only one unique clone in this panel of antibody fragments, termed as VHH-AR185. To investigating the basis of dephosphorylation of RyR2 by VHH-AR185, the binding affinity of VHH-AR185 to RyR2 was measured by surface plasmon resonance. VHHs were purified for these experiments by expressing and secreting from the E. coli cytosol. As shown in Fig. 1D, the affinity (KD) of VHH-AR185 to RyR2 was estimated to be 1.93 nM. The result of affinity studies likely explained the inhibition of RyR2 phosphorylation due to the extremely slow dissociation rate of VHH-AR185 from RyR2.

**Figure 1.**
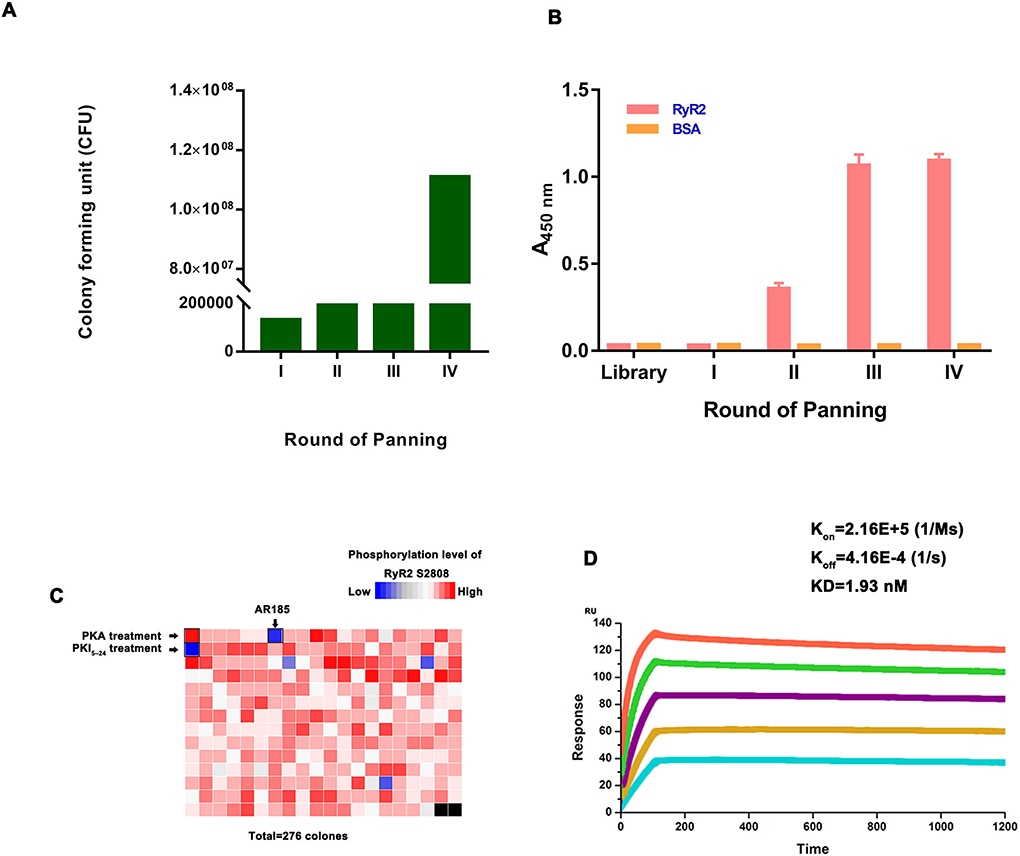
Isolation of RyR2-specific nanobody by phage display. (A) Phage-displayed nanobody fragments were selected against RyR2 by four rounds of panning. A gradual increase in phage titers was detected after each round of panning. (B) Polyclonal phage ELISA from the output phage of each round of panning. Control group used BSA as the irrelevant antigen. (C) Heat map generated from ELISA data of purified RyR2 channels which were phosphorylated in the presence of the PKA. (D) Kinetic analysis of AR185 binding to RyR2 was performed by SPR.

The interaction of VHH-AR185 to RyR2 in the cytoplasm of eukaryotic cells was examined by co-immunoprecipitation experiments. VHH-AR185 and RyR2 were expressed in neonatal cardiomyocytes cells and the lysates of transfected cells were detected. As the result in fig. S1, anti-his antibody was able to efficiently co-precipitate RyR2 from the cells that expressed VHH-AR185-HIS, but could not co-precipitate RyR2 from cells expressing VHH-AR117-HIS. Conversely, anti-RyR2 antibody was able to co-precipitate VHH-AR185-HIS with RyR2, but not VHH-AR185-HIS. This result indicated that the VHH-AR185 could maintain its antigen binding ability in the cytoplasm and fold as a soluble protein.

To identify the epitopes recognized by AR185, phage clones were isolated by panning the PhD.-7 phage display peptide library with AR185. Three rounds of selection were performed, and, at each round, the library was pre-cleared with a control AR177 nanobody. After the third round of panning, the binding of the isolated phage clones to AR185 was determined by ELISA. Sequence analysis of AR185-positive phage clones identified five and six distinct amino acid sequences, respectively (fig. S2A). Alignment of these sequences revealed the consensus motifs DKLAC, which could be aligned with the (2725) DKLAN (2729) sequence located at P2 Domain of RyR2 (fig. S2B).

### Intrabody AR185 rescues cardiac function and reverses remodeling in failing rat myocardium in vivo

We constructed an AAV9 vector containing a VHH-AR185 expression between the two AAV2 inverted terminal repeats and the vector was pseudotyped with a capsid of AAV serotype 9, termed as AAV9.AR185. VHH-AR117 were also constructed as a negative control, termed as AAV9.AR117. To access cardiac expression of VHH, we used the method of adding “self-cleaving” T2A peptide to co-expressed a GFP reporter downstream of VHHs (Fig. 2A). We used the HEK-293 cells expression of different AAV9 particles in vitro and Transmission electron microscope was used to access the AAV9 particles (Fig. 2B). Next, we evaluated the efficiency of gene expression delivered by AAV in vivo. A dosage of AAV9.AR185 particles was delivered to each rat at 1× 10^12^ genome containing particles (gcp), whereas AAV9.AR117 vector was given to the control treated group (n = 5) at the same dosage. After four weeks, we removed all organs from the sacrificed rat, weighed, and assayed treated tissue for fluorescence intensity. Efficiency of gene expression and ability of targeting were evaluated by the ratio of fluorescence intensity to mass of tissue under fluorescence microscope (Fig. 2C and D).

**Figure 2.**
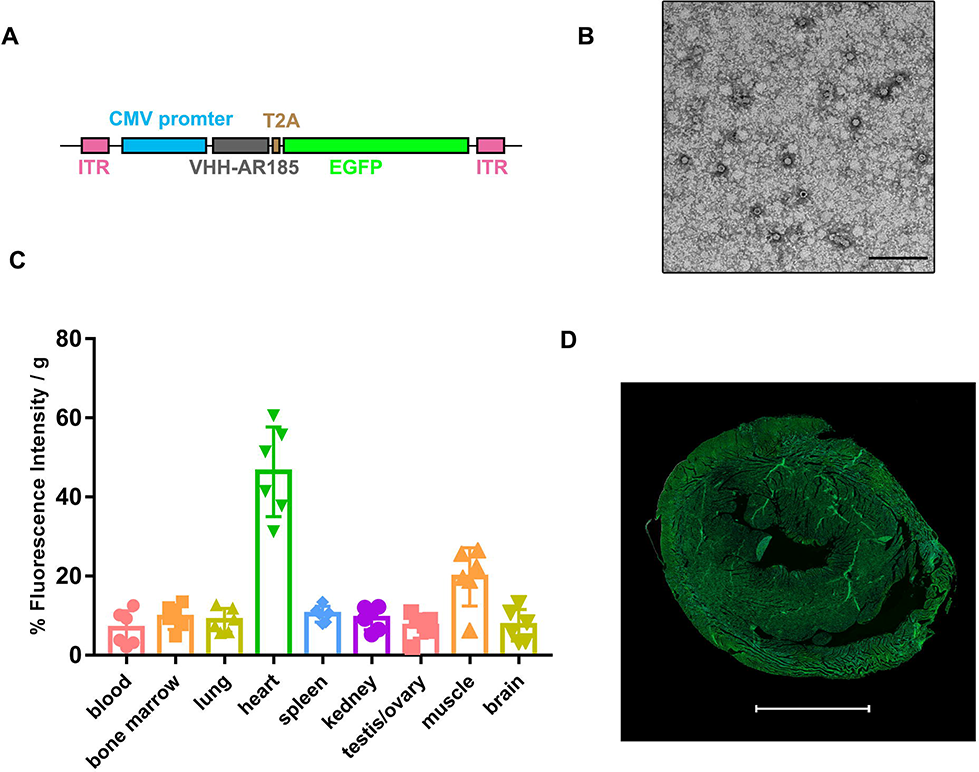
Delivery of a cardiac-specific intracellular nanobody. (A) Schematic diagram of constructs that have the function of expressing nanobody targeting RyR2 along with EGFP by using the T2A sequence. (B) Transmission electron micrographs of AAV9.AR185 particles. The specimens were prepared with negative staining by uranyl acetate. Scale bars = 100 nm. (C) The distribution of AAV9.AR185 expressing in vivo. 4 weeks after intravenous injection of AAV.AR185, SD rats were sacrificed, samples from different tissue were collected and assayed for fluorescence intensity. Data are presented as % FI/g of tissue and displayed as the mean ± SD. (D) Representative fluorescent image of heart that was infected by AAV9.AR185.

To explore the therapeutic potential of VHH, we chose the mode of ischemic heart failure induced by coronary artery ligation for this study. Following the ligation operation, rats were divided into different groups as described in Methods and received control virus (AAV9.AR117), AAV9.AR185 treatment or saline (HF) (n=7-8). The sham-operated animals (Sham) were used as healthy controls. Nine weeks after ligation operation and injection of AAV particles, LV dimensions in the short-axis view was measured by cardiac echo and we also calculated and analyzed the value of ejection fraction and fractional shortening. Our data shows that rats of HF group and AAV9.AR117 group exhibited progressive cardiac dysfunction and LV enlargement, while AAV9.AR185-treated animals showed significant improvement. Moreover, Ejection Fraction and fractional shortening was markedly improved in AAV9.AR185 group compared with HF group and AAV9.AR117 group (Fig. 3A). To determine whether AAV9.AR185 treatment prevented adverse remodeling of the heart after MI, Masson trichrome staining of cardiac sections was performed to measure cardiac fibrosis (Fig. 3B). Whereas there was a significant increase in the development of cardiac fibrosis in Rats of HF group and AAV9.AR117 group after HF, whereas the amount of fibrosis was significant reduced in AAV9.AR185-treated animals. Additionally, HF rat and AAV9.AR117 treated rat had development of a significant increase of heart weight to body weight ratios (HW/BW) after MI compared with sham-operated rat, which is indicative of cardiac remodeling in the context of congestive HF!(Fig. 3C!). In contrast, there was no significant increase in HW/BW ratio after MI in AAV9.AR185-treated rat compared with sham-operated rat. Sarcomeres and mitochondria were the most important index for analysis of ultra-structures of cardiomyocytes from left ventricle that were observed by transmission electron microscopy (Fig. 3D). In the AAV9.AR185 treated and Sham groups, myofilaments were neatly arranged, sarcomeres were intact and Z lines were clear. Conversely, in the HF and AAV9.AR117 groups, MI leaded to disordered arrangement of sarcomeres, dissolution of myofilaments, and frequent vacuoles. In both HF and AAV9.AR117 groups, a lot of mitochondria were swollen and even ruptured, and the separated mitochondrial cristae frequently appeared. The mitochondria in Sham group were well shaped, and the cristae of the mitochondria were obvious and tightly packed. The observations of mitochondria were improved in the AAV9.AR185 treated group compared with AAV9.AR117 treated group. Comprehensively considering the alteration of cardiac function and changes in structure of different groups, the TEM images further support that VHH-AR185 had therapeutic effect in treating heart failure.

**Figure 3.**
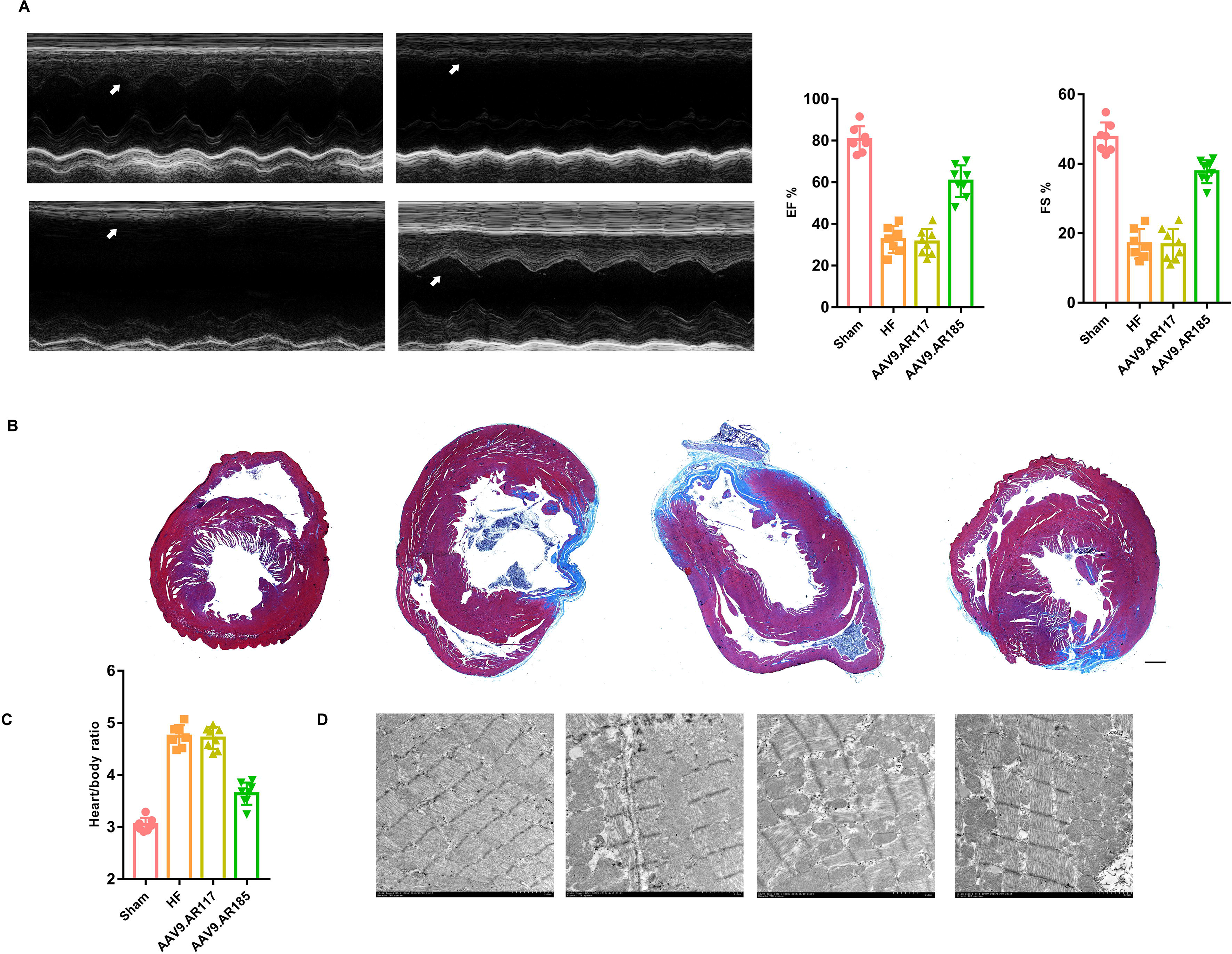
AAV9.AR185 gene therapy rescues cardiac function and reverses remodeling in failing rat myocardium in vivo. (A) Representative wall motion showed by echo cardiograms in different treatment group. Arrows point to septum in all echoes and reduced wall motion appeared in the HF and AAV.AR117 group. In addition, rats in HF and AAV9.AR117 group showed significant reduction of EF and FS (%) compared with Sham and AAV.AR185. **(B)** Representative microstructure of transverse heart sections from four different groups was observed after Masson’s trichrome staining. **(C)** There were significant rises of heart weight to body weight ratio in HF (n =3) and AAV.AR117 treated animals compared with Sham or AAV9.AR185 treated animals. **(F)** The ultrastructure of myocardium acquired by TEM in different treatment groups.

We next accessed the contractile kinetics of isolated LV cardiomyocytes (Table1). When cardiomyocytes were field-stimulated at a frequency of 1 Hz, HF and AAV9.AR117 treated myocytes had significantly slower velocities of shortening and relengthening in than AAV9.AR185 treated myocytes. Fractional shortening of myocytes that were isolated from HF and AAV9.AR117 treated animals also decreased, and time to 50% peak shortening (TPS50%) and time to 50% relengthening (TR50%) became longer. AAV9.AR185 treatment protected cardiomyocytes contractility reserve from the impairment induced by MI. However, only the index of TR50% in myocytes from AAV9.AR185 treated animals returned to a level similar to those of sham operated animals.

### AR185 gene therapy restores cardiomyocyte and SR calcium handling in failing myocardium

We used laser scanning confocal microscopy recorded the fluorescence intensity to measure the sarcoplasmic reticulum Ca^2+^ content of cardiomyocytes from different groups by incubation in the fluorescent dye Fluo-5N/AM. As shown in Fig. 4A, basal sarcoplasmic reticulum Ca^2+^ contents in HF and AAV9.AR117 treated animals were lower than in AAV9.AR185 treated and sham-operated animals.

**Figure 4.**
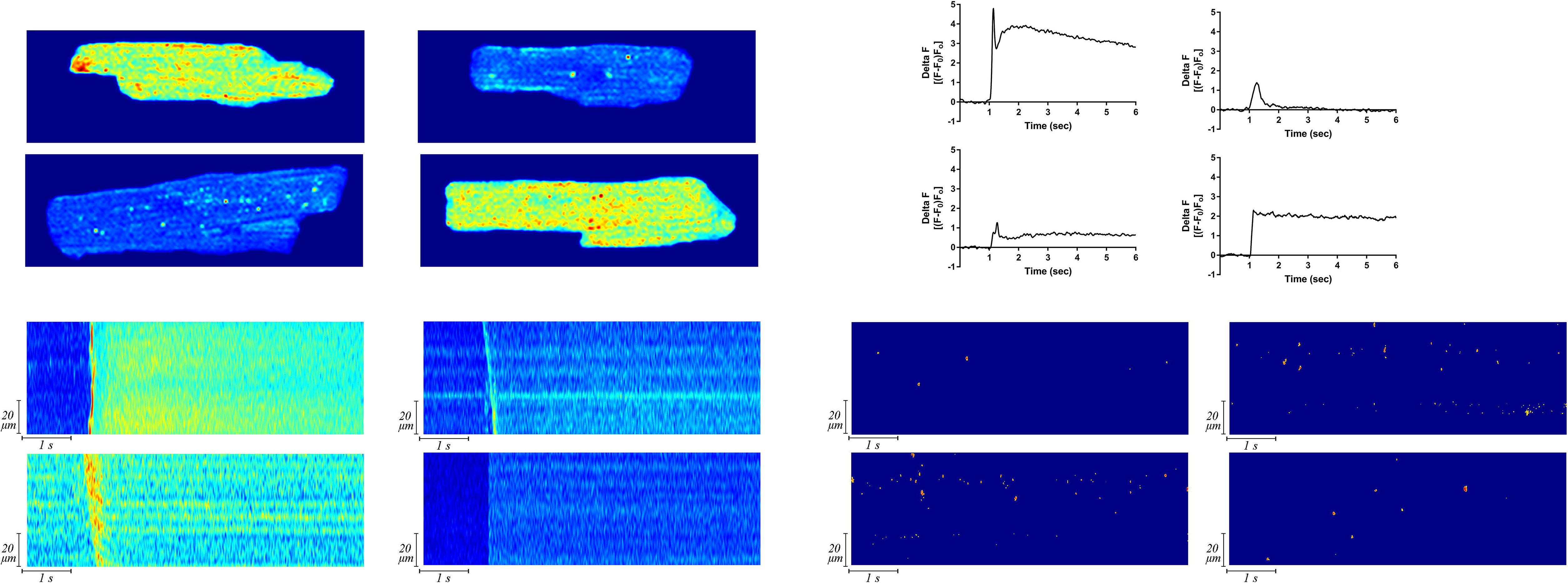
AAV9.AR185 gene therapy restores cardiomyocyte and SR calcium handling in failing myocardium. **(A)** For cardiomyocytes from different treatment group, the fluorescence intensity that reflected sarcoplasmic reticulum Ca^2+^ content was detected by incubating with Fluo-5N/AM. **(B)**. Processed fluorescent images of cardiomyocytes that recorded by line scanning model showed the amplitude of caffeine-induced Ca^2+^ transients in different groups(C) Chart shows Ca2+ transient characteristics of (B). (D) Spontaneous Ca^2+^ sparks appeared during diastole in cardiomyocytes from different treatment group and were showed by the representative processed line-scan images (respectively, n=30 cardiomyocytes from 3 or 4 different hearts were studied for three experiments)

**Figure 5.**
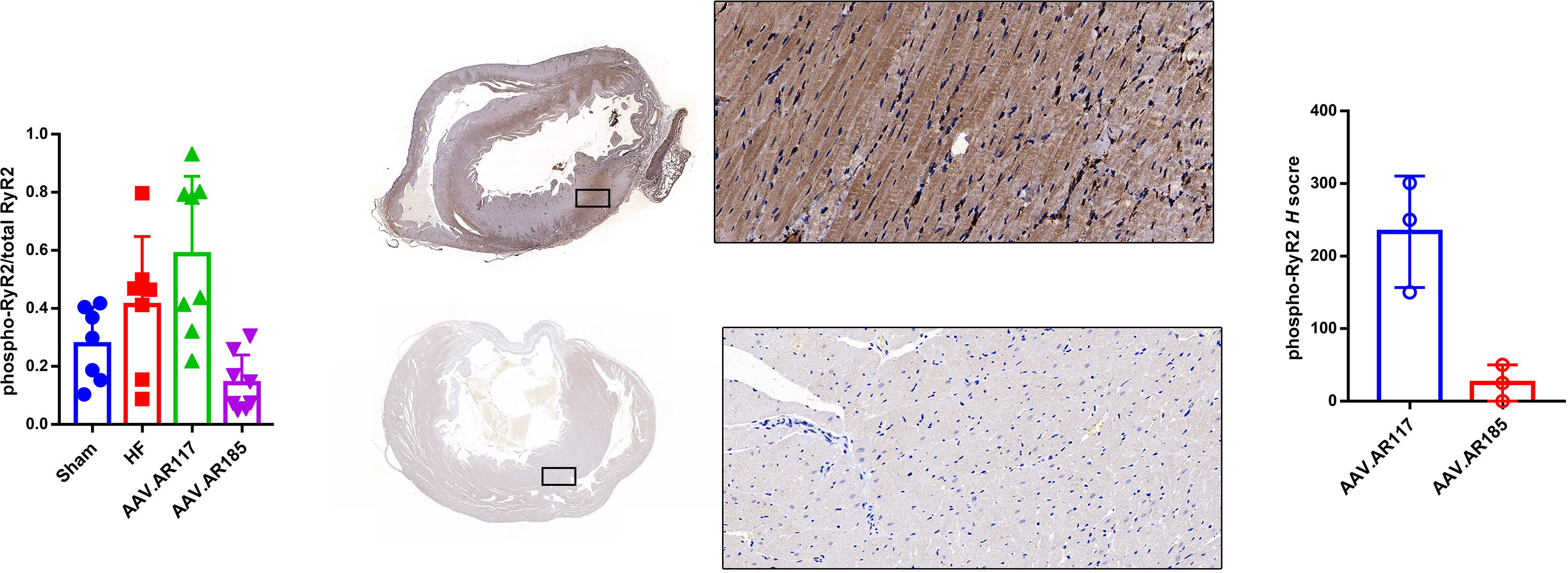
AAV9.AR185 gene therapy inhibits phosphorylation of RyR2 S2808 in failing hearts. (A) Myocardial tissue was harvested, homogenized, and analyzed for pRyR2 (Ser2808) and total RyR2by using ELISA. The statistical significance was determined by using Dunnett’s test. (B) (D) Histologic evaluation of myocardial tissue in different treatment groups. Scale bars = 100 mm. Quantification is expressed as means ± SEM; n = 3. Statistical analysis were done by one-way ANOVA followed by Tukey post-test.a

Additionally, we measured amplitude of calcium transient by incubation in Fluo-4/AM and caffeine perfusion. The representative colorful images in Fig. 4B show line-scan results of evoked Ca^2+^ transients from Shams, HFs, AAV9.AR117s and AAV9.AR185s. When challenged with 20 mM caffeine, less Ca^2+^ was released from the SR of myocytes from AAV9.AR117 group compared with myocytes from AAV9.AR185 treated rats. The results also showed there was significantly reduction of the amplitude of Ca^2+^ transients in the HFs and AR117s compared to that in AR185s. Therefore, the decrease in Ca^2+^ transient amplitude may be the causative factor of the impairment in SR Ca^2+^ load. Rate of Ca^2+^ rise also was significantly slower in HF and AAV9.AR117 myocytes than in AAV9.AR185 treated myocytes (Fig 4C). AAV9.AR185 treatment increased peak of amplitude of evoked Ca^2+^ release and rate of Ca^2+^ rise during Ca^2+^ release.

Table 2 shows representative line-scan images of Ca^2+^ release during the resting stage of cardiomyocytes from Sham (A), HF (B), AAV9.AR117 (C), and AAV9.AR185 animals (D). The data showed that the frequency of Ca^2+^ release was significantly higher and Ca^2+^ sparks occurred frequently in the HF and AAV9.AR117 group compared with the AAV9.AR185 group. The duration of Ca^2+^ sparks in HF and AR117 myocytes were similar to those in Sham and AR185 myocytes, but the Ca^2+^ rise rate of sparks was slower, fluorescence intensity of Ca^2+^ sparks was decreased and T50 decay was longer.

### VHH-AR185 inhibits phosphorylation of RyR2 S2808 in failing hearts

To examine whether AAV9.AR185 treatment results in dephosphorylation of RyR2 and in vivo, cardiomyocyte lysates were further subjected to ELISA analysis, our data shows treatment with AAV9.AR185significantly reduced the level of pRyR2 (S2808) in the cardiomyocytes compared with HF group and AAV9.AR117 treatment (p = 0.0003, Dunnett’s test). Moreover, immunohistochemical analysis of the heart tissues in different treatment group also revealed that an increased accumulation of RyR2 phosphorylation was also observed in the AAV9.AR117 treated group, AAV9.AR185 treatment decreased the level of pRyR2 stain of cells in the myocardium, which indicated that VHH185 has blockage effect of RyR2 phosphorylation. Together, these data demonstrate that AAV9.AR185 treatment leads to inhibition of the RyR2 phosphorylation in vivo. (Fig. 3b1, b2).

## 4 Discussion

In this experiment, we blocked the PKA regulated phosphorylation site(S2808) by exogenously expressing a intracellular antibody specifically binding to RyR2 in cardiomyocytes. After AAV.AR185 treatment, the Ca2+ handing properties of RyR2 in rat hearts isolated form treatment group were very similar to that from Sham group. When the β-adrenergic signaling pathway was activated in vivo, the PKA-regulated site of RyR2 could not bind to phosphate residues, and the phosphorylation of RyR2-S2808 maintained at a low level. At the organic level, AR185 antibody can improve the contractile function of the failing heart, significantly increase the cardiac ejection fraction of heart failure rats, and protect the heart structure from the effects of poor remodeling (Fig 3). At the cellular level, AR185 antibody can reduce the calcium leakage of cardiomyocytes in heart failure rats, maintain the calcium capacity in the sarcoplasmic reticulum of cardiomyocytes, restore calcium homeostasis, and protect the contractility of cardiomyocytes (Fig 4 and Table 1-2). These data is consistent with the previous transgenic animal experiment that down-regulated phosphorylation level of RyR2 S2808 is associated with improved contractility of cardiomyocytes, indicating that S2808 is one of the most important targets in the pathological state of ischemic heart failure.(*11*)

To our knowledge, the evidence for efficiency and safety of using antibodies to treat heart disease was limited. Nanobodies are single domain antibodies consisting of the heavy chain variable domain (VHH) in the camelid family which lacks the light chain. Currently, a variety of nanobodies have entered the clinical research stage.(*29–32*) Compared with traditional antibodies, nanobodies have the advantages of low molecular weight, high affinity, high stability, low immunogenicity and strong penetrability. (*33*) Based on the characteristics of nanobodies and VHH, the use of adeno-associated virus vectors to mediate nanobody treatment of heart failure has great potential. In this study, we successfully expressed AR185 nanobodies which specifically bind to RyR2 in rat cardiomyocytes. Our experimental data demonstrate that intracellular antibody treatment is effective in heart disease rats and does not present a significant safety risk. There are several limitations in our study. We provide evidence that targeting RyR2 with AR185 can be achieved, but rats with heart failure may not fully recapitulate RyR2 in human disease, the experiment data are acquired from a small number of animals, and limited by the shorter observation time, no rats died of heart failure before sacrifice. Furthermore, the mechanisms responsible for therapeutic improvement of AR185 against RyR2 have not been well characterized yet. Hence, these findings will need further validation.

The therapy strategy of AAV9-mediated intracellular antibody which takes advantage of the high specificity and affinity of biomacromolecules can achieve the purpose of specificity regulation of RyR2 and treatment of heart failure. RyR2 is a very promising therapeutic target protein of treatment for heart failure, and intracellular antibody technology in gene therapy is considered as a promising approach. The targets and techniques are worthy of further validation and exploration by researchers.

## Acknowledgments

We thank Y. Bao and Z. Zou (Changzheng Hospital Translational Medicine Center) for generous help of the research. This study was supported by the National Natural Science Foundation of China (grant no. 81773261 and 81602690); Military Medicine Special grant of Second Military Medical University(grant no.2017JS01) a General Financial Grant from the China Postdoctoral Science Foundation (grant no. 2016M593006), and postdoctoral scientific research funds of Second Military Medical University.

